# Desktop iDMS: A stand-alone application of intelligent differential mobility spectrometry (iDMS), a neural network that predicts optimal separation and compensation voltages for field asymmetric differential ion mobility spectrometry

**DOI:** 10.64898/2026.10.02.756329

**Authors:** Thao Nguyen-Tran, Emily Hashimoto-Roth, Xun-Xun Shi, Michael G. Organ, Theodore J. Perkins, Mathieu Lavallée-Adam, Steffany A. L. Bennett

## Abstract

**Summary:** Mass spectrometry resolution of glycosphingolipids requires advanced separation methods. The virtually identical structures of glycolipid epimers cannot be discriminated by routine multiple reaction monitoring-high-performance liquid chromatography-electrospray ionization-tandem mass spectrometry (MRM-LC-ESI-MS/MS). Separation can be achieved using field asymmetric (differential) ion mobility mass spectrometry (FAIMS/DMS). A major bottleneck to MRM-LC-ESI-DMS-MS/MS method development lies in manually optimizing the compound-dependent parameters of separation voltage (SV) and compensation voltage (CoV). To address this challenge, we developed intelligent Differential Mobility Spectrometry (iDMS). iDMS is a supervised deep neural network that evaluates and predicts the optimal ion mobility SV and CoV parameters for any user-requested monoglycosphingolipid from a training set of measured signal intensities composed of 12 lipids (6 stereoisomer pairs). We initially introduced iDMS as a series of python scripts. Here, we share desktop applications of both iDMS and its pre-processing input data assembly modules (DMS Data Extraction Toolkit). Both software assemblies enable users to rapidly deploy a validated machine learning alternative to manual DMS optimization. The applications are user-friendly, operate without need for any technical operating environment, make training dataset assembly quick and simple, and return measured, normalized, and predicted ionograms at every user-interrogated SV, evaluation of separability at each SV and CoV combination, and prediction of optimal resolution parameters to accelerate users’ MRM-LC-ESI-DMS-MS/MS deployment.

**Availability and implementation:** Desktop iDMS and DMS Data Extraction Toolkit are native desktop applications for macOS (Apple Silicon), freely available for download at https://www.neurolipidomics.com/platforms.html under the GNU v3.0 license. The source code is available from GitHub and Zenodo at https://github.com/Neurolipidomics/iDMS, https://zenodo.org/records/22214408, https://github.com/Neurolipidomics/DMSDataExtractionToolkit, and https://ze-nodo.org/records/22777403. Benchmarking metrics are available at https://www.biorxiv.org/content/10.64898/2026.08.26.747394v1.

**Supplementary Information:** Supplementary material is available at *Bioinformatics* online.

## 1 Introduction

Measurement of monoglycosphingolipid stereoisomers at the molecular level is essential to assess target engagement in clinical trials of cause-directed enzyme replacement, substrate reduction, and gene therapy (Ortega, et al., 2022; Ysselstein, et al., 2021); however, routine quantitative high-performance reversed phase liquid chromatography, electrospray ionization, tandem mass spectrometry (MRM-LC-ESI-MS/MS) methods cannot discriminate between glucosyl and galactosyl monoglycosphingolipid stereoisomers (Xu, et al., 2019). For example, homozygous variants in the *GBA1* gene are the genetic determinants of Gaucher Disease (Hruska, et al., 2008), while autosomal dominant-inherited variants account for 10% of Parkinson’s Disease (Cook, et al., 2024). *GBA1* encodes the β-glucocerebrosidase enzyme that hydrolyzes the glucose headgroup from β-glucosylceramides (β-GlcCers) and β-glucosylsphingoid bases (β-GlcSphs) but not the galactose headgroup from β-galactosylceramides (β-GalCers) or β-galactosylsphingoid base (β-GalSphs) stereoisomers (Akiyama, et al., 2020). Conversely, *GALC* encodes β-galactosylceramidase, responsible for removing the galactose but not the glucose headgroup from both simple (mono) and complex glycosphingolipids (Deane, et al., 2011). Routine MRM-LC-ESI-MS/MS methods report combined abundances under the umbrella term, hexosylsphingolipids at every m/z (i.e., *N-*acyl chain length and degree of unsaturation) and thus fail to capture the appropriate substrate under clinical interrogation.

Field asymmetric ion mobility mass spectrometry (FAIMS) also known as differential ion mobility spectrometry (DMS) can resolve β-Glc and β-Gal-containing sphingolipids in MRM-Lunique to each user’s DMSC-ESI-DMS-MS/MS methods when used as an orthogonal separation to LC (Xu, et al., 2019). The interaction of glycosphingolipid ions with the neutral modifier gas molecules in the DMS cell promotes formation of ion clusters according to their sugar head-group. These discrete ion clusters can then be selectively introduced into the MS, first by diverting the clusters away from the MS inlet through the application of an alternating current (known as separation voltage or SV) and then sequentially redirecting either β-Glc or β-Gal stereoisomer ions selectively back into the MS by the application of a direct current known as compensation voltage (CoV). The bottleneck to deploying quantitative MRM-LC-ESI-DMS-MS/MS methods is the time-consuming process of establishing the monoglycosphingolipid-dependent optimal SV and CoV combinations. Importantly, these parameters can only be empirically established using lipid standards. Without standards, method development is not possible.

To address this problem, we developed intelligent Differential Mobility Spectrometry (iDMS) (Nguyen-Tran, et al., 2026). iDMS is a supervised deep neural network that predicts the optimal ion mobility parameters required to resolve any monoglycosphingolipid stereoisomer pair according to their sugar headgroup, *N*-acyl chain length, and *N*-acyl degree of unsaturation. Resolution parameters for species that do not have commercial standards can be predicted by learning the non-linear relationships between SV, CoV, and monoglycosphingolipid structure unique to each user’s DMS platform. iDMS requires a minimal training dataset of 12 lipids (6 stereoisomer pairs) to predict parameters for any β-Glc and β-Gal Cer. iDMS was released as two Python scripts and two Jupyter notebooks (Nguyen-Tran, et al., 2026). To more rapidly field-deploy this machine learning solution to users without need for technical environmental or dependency setup, we introduce here a user-friendly iDMS desktop application supported by a DMS Data Extraction toolkit to batch extract users’ data from .jdx or .mzml acquisition files into the required iDMS input format for training. Desktop iDMS returns both training and *in silico* predicted data as ionograms according to the user’s FAIMS/DMS-specific platform, determines resolution at each theoretical combination, and predicts the optimal SV and CoV combinations for deployment in the users MRM-LC-ESI-DMS-MS/MS method of choice.

## 2 Implementation

DMS Data Extraction Toolkit and iDMS are implemented in Python 3.12 as two desktop applications. Each application is a structured orchestration of the individual iDMS Python scripts providing a user-friendly option for running the pipeline from data acquisition, modelling, extracting the training features, executing the neural net, and evaluating predictions (Fig 1). iDMS requires two input .csv files corresponding to paired lipid isomers with columns for CoV, intensity, SV, chain length, unsaturation unit, isomer identity, sphingoid backbone, and lipid-species annotation. These inputs are parsed and validated for shared lipid species in both files (i.e., both isomer files contain empirical data for lipids with the same *N*-acyl chains) to provide a controlled and reproducible entry point to the downstream prediction modelling. Lastly, iDMS requires the user to input the lipids for which they would like to predict the optimal SV and CoV values. This may be typed directly into the application or uploaded a .csv file.

**Fig. 1.**
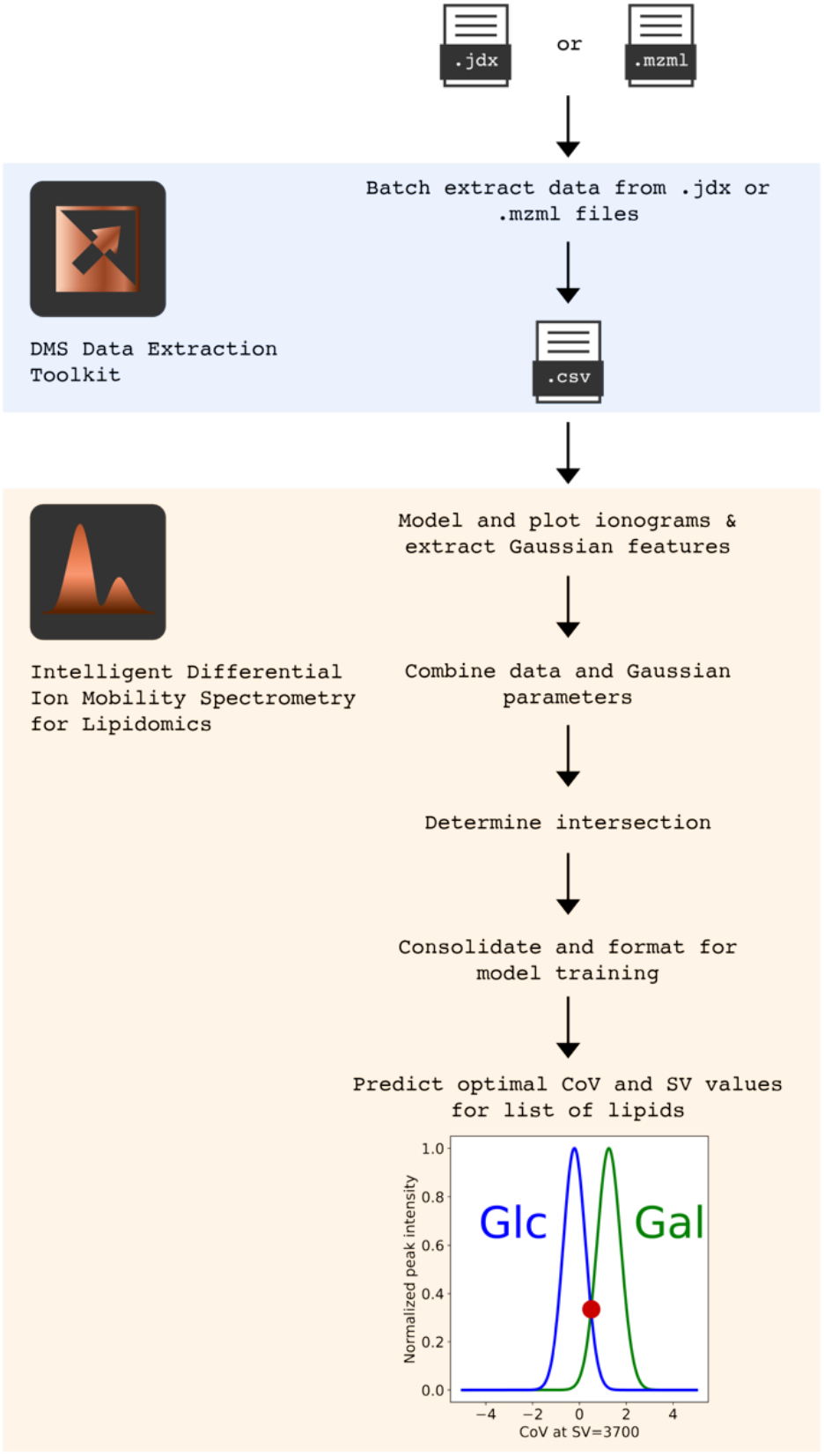
Overview of the iDMS pre-processing, training, and prediction workflow. Outputs are .csv files and .pngs for normalized signals from experiments, gaussian parameters derived from training data, and the intersection point of the isomers determining separability, with all respective plots. Examples of the iDMS results (files and plots) are provided in Supplementary Figures 1-6.

Because iDMS requires specifically formatted input data, we created the DMS Data Extraction Toolkit, combined as one application called DMSExtract. This toolkit can be used to batch extract and format the data from a collection of .jdx or .mzml files to generate the .csv input files.

To run the toolkit, users must provide via an easy-to-use UI:

- CoV range and increment
- Glyco-headgroup
- Sphingoid backbone
- Parent directory containing experimental sub-directories for individual lipid species

The DMS Data Extraction Toolkit automatically detects the experimental sub-directories and prompts the user to enter the *N*-acyl chain length and unsaturation unit of each experiment. Supplementary Figure 1 presents the processing and output of DMS Data Extraction Toolkit.

To streamline the iDMS workflow, the application combines both iDMS modules. The original iDMS scripts were organized in two modules. Module 1 estimates and models the Gaussian parameters of the input lipid isomers. Module 2 executes the supervised deep neural network and assesses separability of the predicted CoV and SV values. To accelerate this process, the iDMS desktop application presented here integrates both modules into a single consolidated pipeline. A key implementation feature is the explicit management of output directories and reusable intermediate files, wherein each run stores its outputs in a defined directory structure. Separate sub-directories are generated for the Gaussian parametric modelling and supervised model predictions. The application also creates a .json file (“lipid_data.json”) containing the isomer labels, sphingoid backbone, and list of shared lipids to preserve the information needed to identify the datasets used in a run. This .json file is leveraged in the application’s “skip mode” for cases when the user wants to predict for new lipids without retraining their model. This feature allows the user flexibility and a means to save time during their analyses not available in the original modular scripts.

## Conclusion

iDMS and DMS Data Extraction Toolkit are free open-source field-ready desktop applications. These desktop application implementations are designed as practical analysis applications as compared to the methodological protype provided by the original scripts. By combining standardized input requirements, combined modular execution, and structured output, iDMS provides a reproducible software framework for rapidly deploying this machine learning alternative to empirical DMS parameter optimization and prediction for lipids lacking commercial standards. Importantly, these applications enable users to establish, for the first time, the FAIMS/DMS parameters required to resolve stereoisomers for which standards are not available for compound-dependent optimization, thereby expanding capacity of clinical glycosphingolipidomics. User guides and sample files for both the DMS Data Extraction Toolkit (DMSExtract) and iDMS are available for download at https://www.neurolipidomics.com/platforms.html, from the File menu in each application, and as part of Supplementary Data 7_User Guide: Understanding the DMS Data Extraction Toolkit Application Interface and 8_User Guide: Understanding the iDMS Interface.

## Data availability

All software, resources, and code associated with iDMS and the DMS Data Extraction Toolkit are available via GitHub and Zenodo cited in the Availability and Implementation section of the abstract. Benchmarking data and methodology are available in (Nguyen-Tran, et al., 2026)

## References

Akiyama, H., et al. Glucocerebrosidases catalyze a transgalactosylation reaction that yields a newly-identified brain sterol metabolite, galactosylated cholesterol. J Biol Chem 2020;295(16):5257–5277.

Cook, L., et al. Parkinson’s disease variant detection and disclosure: PD GENEration, a North American study. Brain 2024;147(8):2668–2679.

Deane, J.E., et al. Insights into Krabbe disease from structures of galactocerebrosidase. Proc Natl Acad Sci U S A 2011;108(37):15169–15173.

Hruska, K.S., et al. Gaucher disease: mutation and polymorphism spectrum in the glucocerebrosidase gene (GBA). Human mutation 2008;29(5):567–583.

Nguyen-Tran, T., et al. Intelligent differential ion mobility spectrometry (iDMS): A deep neural network that predicts optimal space-resolved ion mobility parameters for isomeric monoglycosphingolipids. bioRxiv 2026:2026.2008.2026.747394.

Ortega, R.A., et al. Assessment of Glucocerebrosidase Enzyme Activity in Parkinson Disease Using Multiple Approaches. Movement disorders : official journal of the Movement Disorder Society 2022;37(3):655–656.

Xu, H., et al. DMS as an orthogonal separation to LC/ESI/MS/MS for quantifying isomeric cerebrosides in plasma and cerebrospinal fluid. J Lipid Res 2019;60(1):200–211.

Ysselstein, D., et al. Evaluation of Strategies for Measuring Lysosomal Glucocerebrosidase Activity. Movement Disorders 2021;36(12):2719–2730.

