## Supplementary Data for "Desktop iDMS: A stand-alone application of intelligent differential mobility spectrometry (iDMS), a neural network that predicts optimal separation and compensation voltages for field asymmetric differential ion mobility spectrometry"

### 1. DMSExtract (DMSExtractionToolkit) output

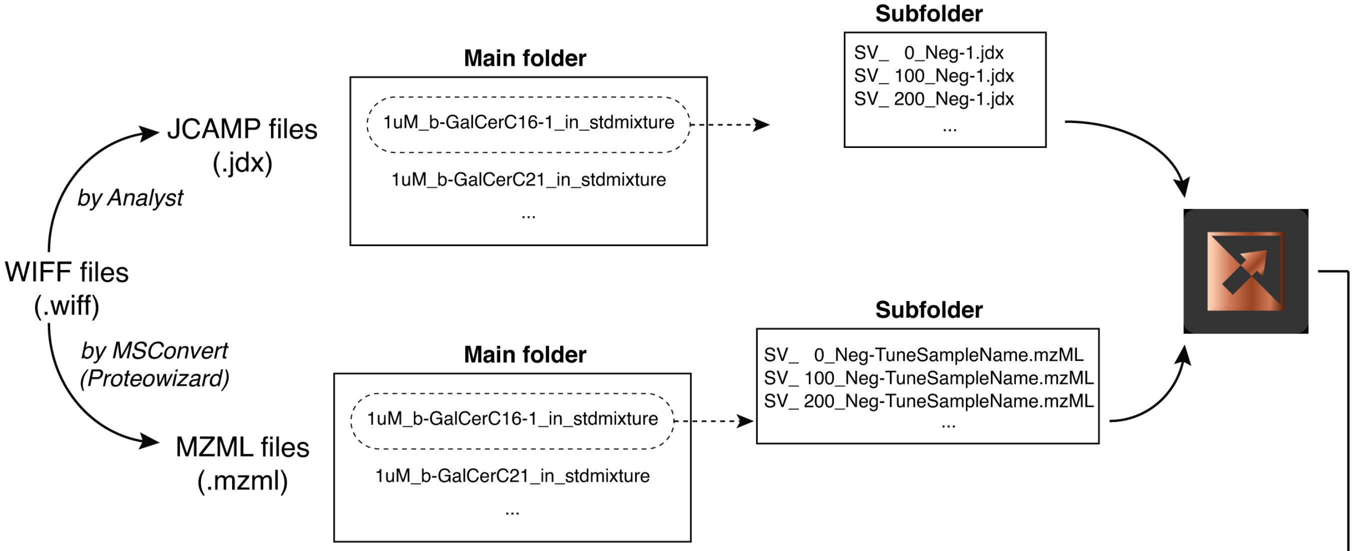

**a Output from JCAMP (.jdx) files**

| COV | Intensity | SV | chainLength | unsaturationUnit | isomer | sphingoidBackbone | lipidSpecies | acquisitionDate | Instrument |
|---|---|---|---|---|---|---|---|---|---|
| -10 | 0 | 0 | 16 | 1 | Gal | d18:1 | 16:1 | 15-09-2020 | SCIEX |
| -9.8 | 0 | 0 | 16 | 1 | Gal | d18:1 | 16:1 | 15-09-2020 | SCIEX |
| … | … | … | … | … | … | … | … | … | … |

**b Output from .mzml files**

| COV | Intensity | SV | chainLength | unsaturationUnit | isomer | sphingoidBackbone | lipidSpecies | acquisitionDate | Instrument |
|---|---|---|---|---|---|---|---|---|---|
| -10 | 0 | 0 | 16 | 1 | Gal | d18:1 | 16:1 | 15-09-2020 | QTRAP_5500_sn_# |
| -9.8 | 0 | 0 | 16 | 1 | Gal | d18:1 | 16:1 | 15-09-2020 | QTRAP_5500_sn_# |
| … | … | … | … | … | … | … | … | … | … |

**Figure S1. Preprocessing Input Dataset Assembly.** DMSExtract batch extracts user's empirical data from .jdx or .mzml acquisition files into the iDMS input format required to generate training datasets. **a)** Formatted input data from .jdx files include acquisition data and instrument metadata in addition to all information needed for iDMS to produce training datasets. **b)** Formatted input data generated from .mzml files include the same data as well as instrument serial number where collected.

### 2. iDMS Module 1 Parametric Modelling: Measured Ionograms

a

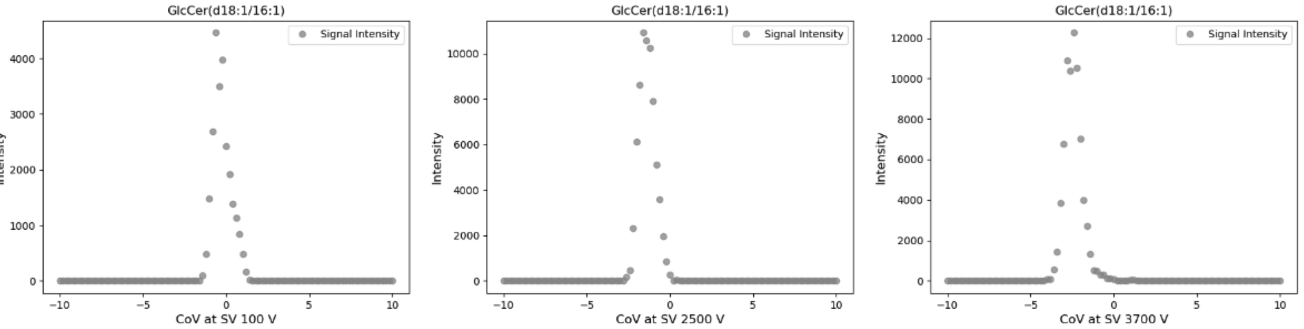

extract_Glc_MaxCov.csv

| COV | Intensity | SV | chainLength | unsaturationUnit | isomer | sphingoidBackbone | lipidSpecies |
|---|---|---|---|---|---|---|---|
| -0.6 | 4460 | 100 | 16 | 1 | Glc | d18:1 | 16:1 |
| -1.6 | 10920 | 2500 | 16 | 1 | Glc | d18:1 | 16:1 |
| -2.4 | 12260 | 3700 | 16 | 1 | Glc | d18:1 | 16:1 |

b

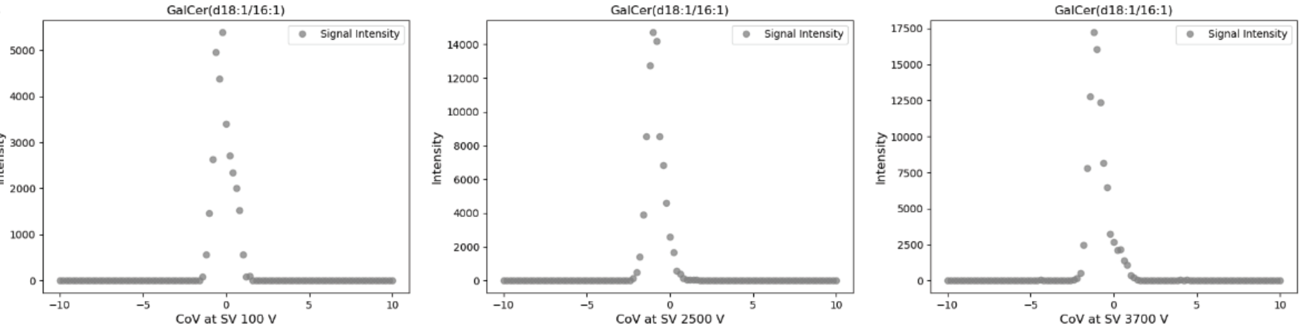

extract_Gal_MaxCov.csv

| COV | Intensity | SV | chainLength | unsaturationUnit | isomer | sphingoidBackbone | lipidSpecies |
|---|---|---|---|---|---|---|---|
| -0.2 | 5380 | 100 | 16 | 1 | Gal | d18:1 | 16:1 |
| -1 | 14700 | 2500 | 16 | 1 | Gal | d18:1 | 16:1 |
| -1.2 | 17220 | 3700 | 16 | 1 | Gal | d18:1 | 16:1 |

**Figure S2. iDMS returns all measured signal intensities in the user's input dataset as ionograms, plotting intensity for each SV as a function of CoV, and extracting the CoV at maximal signal intensity.** Plots are provided in Module1_Outputs/ExtractIonograms as .png files and data returned two .csv files for both epimers. **a)** extract_Glc_MaxCov and **b)** extract_Gal_MaxCov

### 3. iDMS Module 1 Parametric Modelling: Normalized Ionograms

a

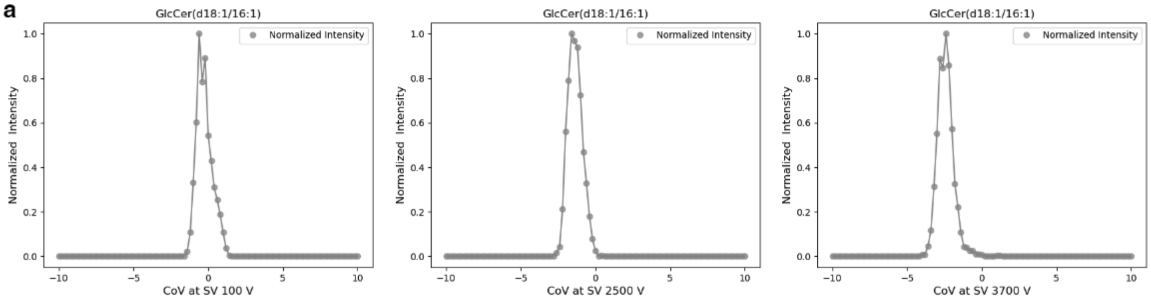

norm_Glc.csv

| COV | Intensity | SV | chainLength | unsaturationUnit | isomer | sphingoidBackbone | lipidSpecies | NormPeakIntensity |
|---|---|---|---|---|---|---|---|---|
| -10 | 0 | 100 | 16 | 1 | Glc | d18:1 | 16:1 | 0 |
| … | … | … | … | … | … | … | … | … |
| -1.4 | 100 | 100 | 16 | 1 | Glc | d18:1 | 16:1 | 0.022421525 |
| -1.2 | 480 | 100 | 16 | 1 | Glc | d18:1 | 16:1 | 0.107623318 |
| -1 | 1480 | 100 | 16 | 1 | Glc | d18:1 | 16:1 | 0.331838565 |
| -0.8 | 2680 | 100 | 16 | 1 | Glc | d18:1 | 16:1 | 0.600896861 |
| -0.6 | 4460 | 100 | 16 | 1 | Glc | d18:1 | 16:1 | 1 |
| -0.4 | 3500 | 100 | 16 | 1 | Glc | d18:1 | 16:1 | 0.784753363 |
| -0.2 | 3980 | 100 | 16 | 1 | Glc | d18:1 | 16:1 | 0.892376682 |
| 0 | 2420 | 100 | 16 | 1 | Glc | d18:1 | 16:1 | 0.542600897 |
| 0.2 | 1920 | 100 | 16 | 1 | Glc | d18:1 | 16:1 | 0.430493274 |
| 0.4 | 1380 | 100 | 16 | 1 | Glc | d18:1 | 16:1 | 0.30941704 |
| 0.6 | 1140 | 100 | 16 | 1 | Glc | d18:1 | 16:1 | 0.255605381 |
| 0.8 | 840 | 100 | 16 | 1 | Glc | d18:1 | 16:1 | 0.188340807 |
| 1 | 480 | 100 | 16 | 1 | Glc | d18:1 | 16:1 | 0.107623318 |
| 1.2 | 160 | 100 | 16 | 1 | Glc | d18:1 | 16:1 | 0.035874439 |
| 1.4 | 20 | 100 | 16 | 1 | Glc | d18:1 | 16:1 | 0.004484305 |
| … | … | … | … | … | … | … | … | … |

b

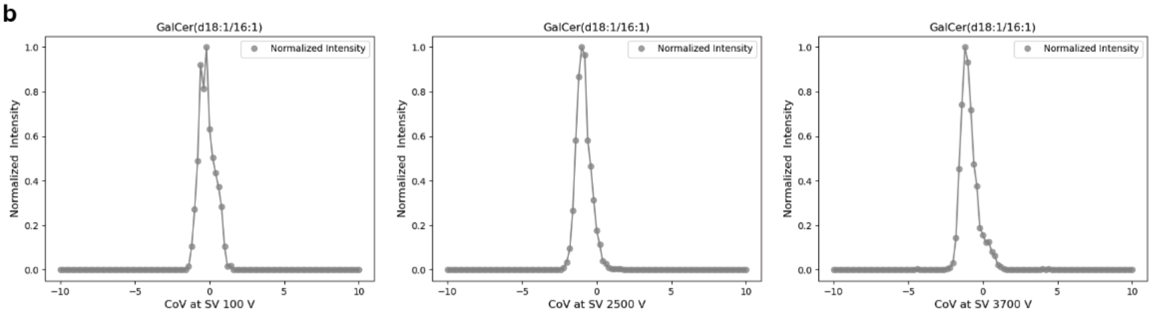

norm_Gal.csv

| COV | Intensity | SV | chainLength | unsaturationUnit | isomer | sphingoidBackbone | lipidSpecies | NormPeakIntensity |
|---|---|---|---|---|---|---|---|---|
| -10 | 0 | 100 | 16 | 1 | Gal | d18:1 | 16:1 | 0 |
| … | … | … | … | … | … | … | … | … |
| -1.4 | 80 | 100 | 16 | 1 | Gal | d18:1 | 16:1 | 0.014869888 |
| -1.2 | 560 | 100 | 16 | 1 | Gal | d18:1 | 16:1 | 0.104089219 |
| -1 | 1460 | 100 | 16 | 1 | Gal | d18:1 | 16:1 | 0.271375465 |
| -0.8 | 2640 | 100 | 16 | 1 | Gal | d18:1 | 16:1 | 0.49070632 |
| -0.6 | 4960 | 100 | 16 | 1 | Gal | d18:1 | 16:1 | 0.921933086 |
| -0.4 | 4380 | 100 | 16 | 1 | Gal | d18:1 | 16:1 | 0.814126394 |
| -0.2 | 5380 | 100 | 16 | 1 | Gal | d18:1 | 16:1 | 1 |
| 0 | 3400 | 100 | 16 | 1 | Gal | d18:1 | 16:1 | 0.63197026 |
| 0.2 | 2720 | 100 | 16 | 1 | Gal | d18:1 | 16:1 | 0.505576208 |
| 0.4 | 2340 | 100 | 16 | 1 | Gal | d18:1 | 16:1 | 0.434944238 |
| 0.6 | 2000 | 100 | 16 | 1 | Gal | d18:1 | 16:1 | 0.371747212 |
| 0.8 | 1520 | 100 | 16 | 1 | Gal | d18:1 | 16:1 | 0.282527881 |
| 1 | 560 | 100 | 16 | 1 | Gal | d18:1 | 16:1 | 0.104089219 |
| 1.2 | 80 | 100 | 16 | 1 | Gal | d18:1 | 16:1 | 0.014869888 |
| 1.4 | 100 | 100 | 16 | 1 | Gal | d18:1 | 16:1 | 0.018587361 |
| … | … | … | … | … | … | … | … | … |

**Figure S3. iDMS normalizes measured signal intensities at each SV to the maximum intensity across all CoVs at that SV, and outputs plots of normalized ionograms and data of measured and normalized intensities.** Data are provided in a Module1_Outputs /NormIonograms as .png plots and as two .csv files. **a)** norm_Glc.csv and **b)** norm_Gal.csv

### 4. iDMS Module 1 Parametric Modelling: Gaussian Fits

a

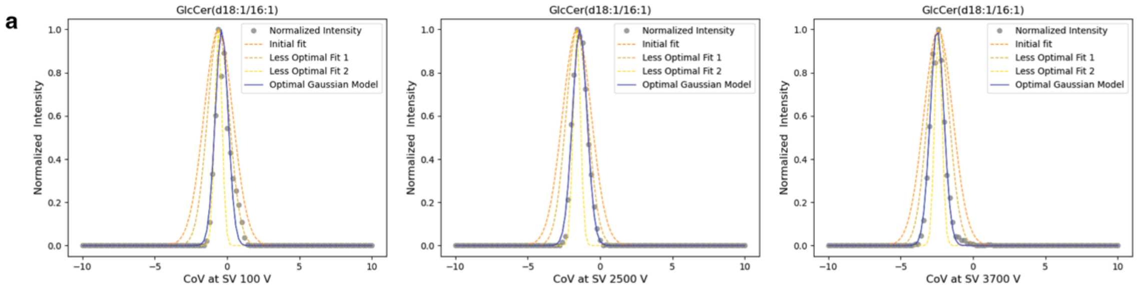

norm_GlcParameters.csv

| COV | Intensity | SV | chainLength | unsaturationUnit | isomer | sphingoidBackbone | lipidSpecies | NormPeakIntensity | Empirical mu | Glc_Gaussian_mu | Glc_Gaussian_sigma | Glc_Rsquared |
|---|---|---|---|---|---|---|---|---|---|---|---|---|
| -0.6 | 4460 | 100 | 16 | 1 | Glc | d18:1 | 16:1 | 1 | -0.6 | -0.4 | 0.445 | 0.9486 |
| -1.6 | 10920 | 2500 | 16 | 1 | Glc | d18:1 | 16:1 | 1 | -1.6 | -1.4 | 0.504 | 0.9914 |
| -2.4 | 12260 | 3700 | 16 | 1 | Glc | d18:1 | 16:1 | 1 | -2.4 | -2.5 | 0.476 | 0.9913 |

b

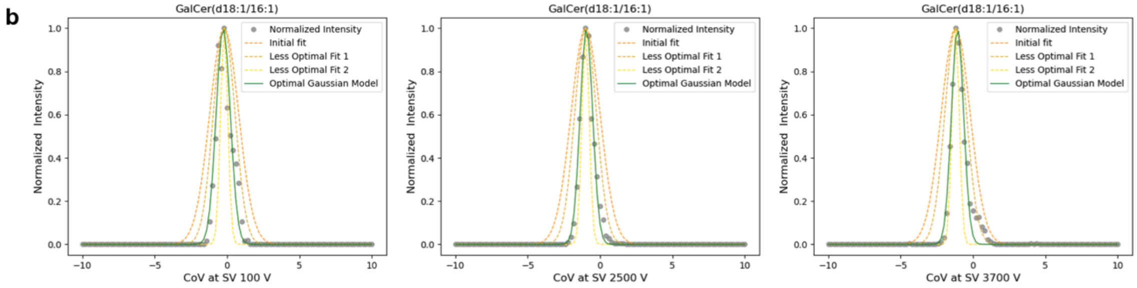

norm_GalParameters.csv

| COV | Intensity | SV | chainLength | unsaturationUnit | isomer | sphingoidBackbone | lipidSpecies | NormPeakIntensity | Empirical mu | Gal_Gaussian_mu | Gal_Gaussian_sigma | Gal_Rsquared |
|---|---|---|---|---|---|---|---|---|---|---|---|---|
| -0.2 | 5380 | 100 | 16 | 1 | Gal | d18:1 | 16:1 | 1 | -0.2 | -0.3 | 0.490 | 0.9462 |
| -1 | 14700 | 2500 | 16 | 1 | Gal | d18:1 | 16:1 | 1 | -1 | -0.9 | 0.434 | 0.9854 |
| -1.2 | 17220 | 3700 | 16 | 1 | Gal | d18:1 | 16:1 | 1 | -1.2 | -1.1 | 0.421 | 0.9713 |

**Figure S4. iDMS fits the normalized intensities to the best-fit Gaussian function deriving CoV $\mu$ and $\sigma$ (at each SV) and returns both ionograms and list data.** Data are returned for both stereoisomers in Module1_Outputs /GaussianIntersectPlots as .png ionograms and as three .csv files. **a)** norm_GlcParameters.csv, **b)** norm_GalParameters.csv and meanAllsigma.csv (not shown).

### 5. iDMS Module 1 Parametric Modelling: Intersection Points

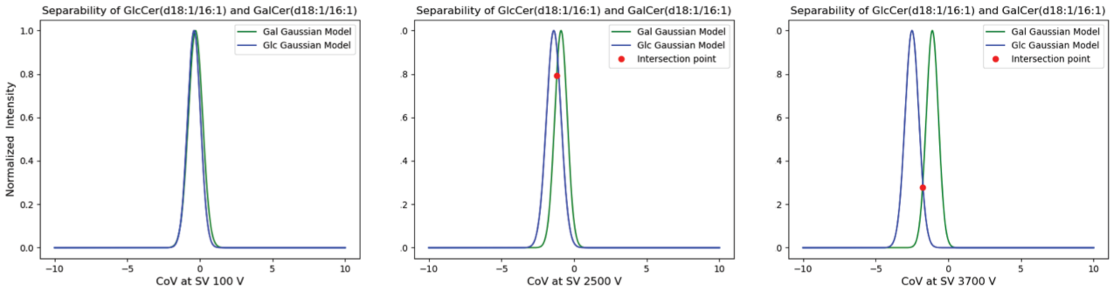

CombinedAnalysisApp.csv
Intersected gauss.csv (includes separation classification)

| lipidSpecies | SV | chainLength | unsaturationUnit | sphingoidBackbone | mu_Gal | sigma_Gal | Intensity_at_mu_Gal | mu_Glc | sigma_Glc | Intensity_at_mu_Glc | Valley CoV | Intersection point | Norm Intensity at Intersection | Sep or InSep |
|---|---|---|---|---|---|---|---|---|---|---|---|---|---|---|
| 16:1 | 100 | 16 | 1 | d18:1 | -0.3 | 0.490 | 5380 | -0.4 | 0.445 | 4460 | | | | Inseparable |
| 16:1 | 2500 | 16 | 1 | d18:1 | -0.9 | 0.434 | 14700 | -1.4 | 0.504 | 10920 | | -1.2 | 0.79212302 | Inseparable |
| 16:1 | 3700 | 16 | 1 | d18:1 | -1.1 | 0.421 | 17220 | -2.5 | 0.476 | 12260 | -1.8 | -1.8 | 0.277181838 | Separable |

**Figure S5. iDMS plots both stereoisomer Gaussian fits, determines their intersection points, and classifies separability**. Separability is defined as the SV and CoV combinations at which the intersection point between the two intensity distributions is <0.5 of the normalized signal intensity. Data are returned for both stereoisomers in Module1_Outputs /Glc_Gaussian Plots or Gal_Gaussian Plots as .png ionograms and as two .csv files, CombinedAnalysisApp.csv and Intersected_gauss.csv that includes classification of separability.

### 6. iDMS Module 2: Prediction

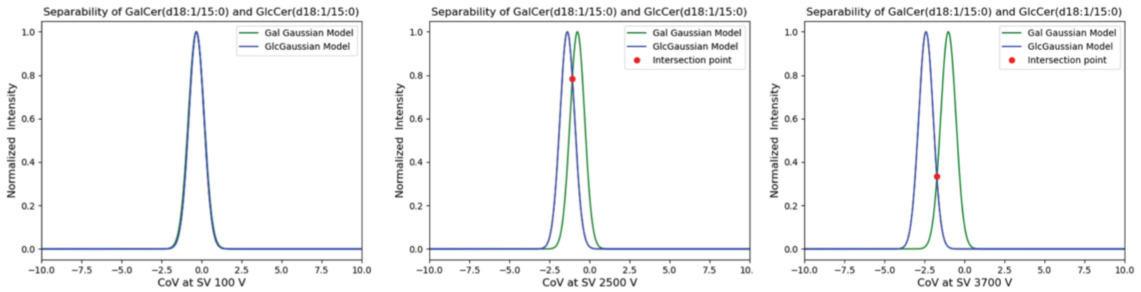

PredictionResultApp.csv

| lipidSpecies | SV | chainLength | unsaturationUnit | mu_Gal | mu_Glc | sigma_Gal | sigma_Glc | Valley CoV | Intersection point | Norm Intensity at Intersection | Sep or InSep |
|---|---|---|---|---|---|---|---|---|---|---|---|
| 15:0 | 100 | 15 | 0 | -0.329 | -0.320 | 0.497 | 0.474 | | | | Inseparable |
| 15:0 | 2500 | 15 | 0 | -0.759 | -1.388 | 0.455 | 0.459 | | -1.075 | 0.785 | Inseparable |
| 15:0 | 3700 | 15 | 0 | -1.008 | -2.403 | 0.478 | 0.457 | -1.714 | -1.714 | 0.336 | Separable |

**Figure S6. iDMS predicts the Gaussian distributions of any stereoisomer pair queried by the user and their intersection points and classifies each SV/CoVμ combination as separable or inseparable**. Data are returned for both stereoisomers in Module2_Outputs /PredictedSetPlots as .png ionograms and as a PredictionResultApp.csv file.

**Neurolipidomics Laboratory**
**India Taylor Lipidomics Research Platform**
Contact: Dr. Steffany Bennett, Thao Nguyen-Tran, Emily Hashimoto-Roth


### Understanding the DMS Data Extraction Toolkit Application Interface:

Specify the CoV parameters, lipid properties, input, and output directories to extract data from a collection of .mzml or .jdx files, saved to a single .csv file.

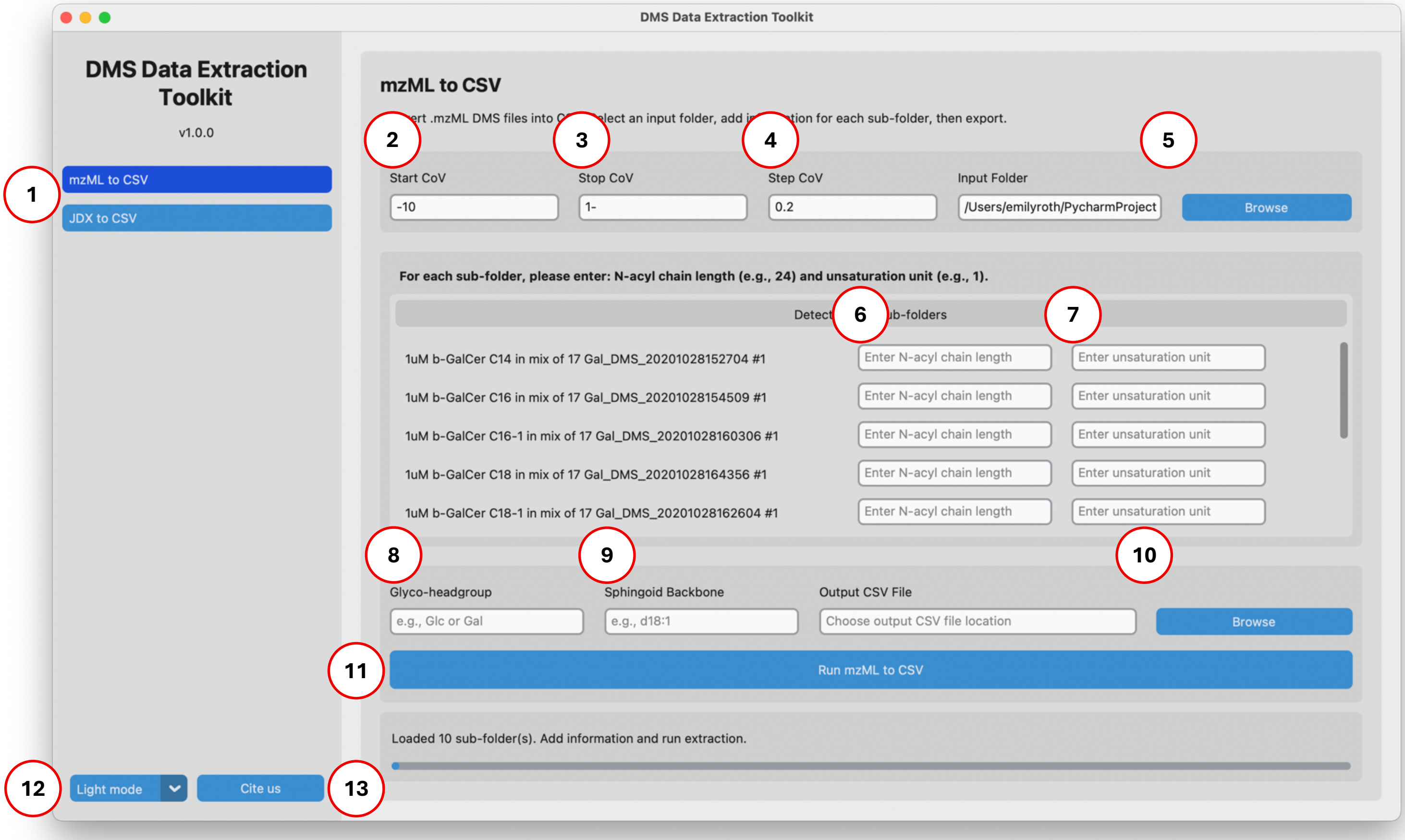

1. **Select tool:** Select the option to extract from either .mzml or .jdx files.
2. **Start CoV:** Input the start value of the CoV range.
3. **Stop CoV:** Input the stop value of the CoV range.
4. **Step CoV:** Input the step by which the CoV increases.
5. **Input folder & browse button:** Select the directory where the input sub-directories are located.
6. **Input *N*-acyl chain length:** For each sub-directory, specify the given *N*-acyl chain length as a single number (i.e., 14)
7. **Input unsaturation unit:** For each sub-directory, specify the given unsaturation unit as a number (i.e., 0)
8. **Glyco-headgroup:** Specify which sugar headgroup is relevant to the set of input experimental runs uploaded

**Neurolipidomics Laboratory**
**India Taylor Lipidomics Research Platform**
Contact: Dr. Steffany Bennett, Thao Nguyen-Tran, Emily Hashimoto-Roth


9. **Sphingoid backbone:** Specify the sphingoid backbone relevant to the set of input experimental runs. Use the following nomenclature (e.g., d18:0, d18:1, d18:2)
10. **Output browse:** Select the directory where DMS Extract will save the extracted data (.csv).
11. **Run:** Start the extraction pipeline.
12. **Theme selector:** Change the application appearance between light (default) and dark mode.
13. **Cite us:** Open the citation information for DMS Extract in a browser.

**Understanding the DMS Extract Application Output:**

DMS Extract outputs a single .csv file formatted for use as sample data input for iDMS.

**Neurolipidomics Laboratory**
**India Taylor Lipidomics Research Platform**
Contact: Dr. Steffany Bennett, Thao Nguyen-Tran, Emily Hashimoto-Roth


### Understanding the iDMS Application Interface:

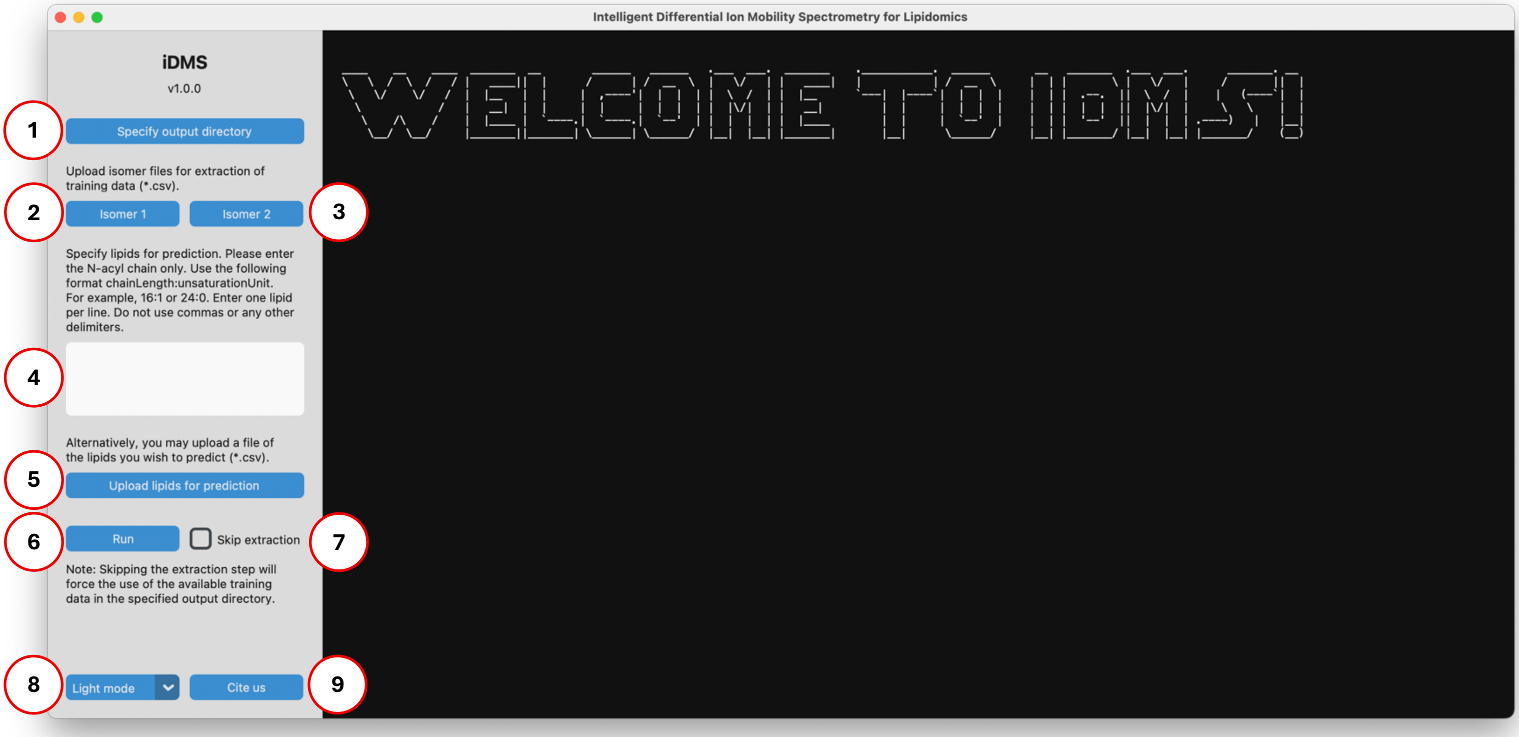

1. **Specify output directory:** Select the directory where iDMS will save extracted (training) data, predictions, and other output files.
2. **Isomer 1:** Upload the first isomer data file (.csv).
3. **Isomer 2:** Upload the second isomer data file (.csv).
4. **Lipid input field:** Enter the lipids you wish to predict. Provide only the *N*-acyl chain information using the format chainLength:unsaturationUnit (e.g.,14:0). Provide only one lipid per line. Do not include additional punctation (e.g., commas).

   Good example: 14:0
   16:0
   16:1

   Bad example 1: 14:0,
   16:0,
   16:1

   Bad example 2: 14:0,16:0,16:1

5. **Upload lipids for prediction:** Alternatively, upload a file (.csv) containing the lipids for prediction. If uploading a list, do not enter the lipids manually. Format must match the format above with one lipid per line.
6. **Run:** Start the analysis pipeline.
7. **Skip extraction:** Select this option to bypass the data extraction step and to instead use existing extracted data (i.e., data extracted from your sample isomer datasets from the last full iDMS run made. If you close iDMS, you must re-extract your data).

8. **Theme selector:** Change the application appearance between light (default) and dark mode.
9. **Cite us:** Opens the iDMS website with citation information in a browser.

**Understanding the iDMS Application Output:**

iDMS generates two sub-directories and a .json file per run. Each sub-folder corresponds to a high-level analysis pipeline step:

- **Module 1 sub-directory:** Extracting and consolidating data from the two input isomer files.
- **Module 2 sub-directory:** Formatting consolidated data for neural network model training and predicting instrument parameters of queried lipids.

The .json file summarizes the input data of a given run and is critical, along with Module 1 results, for running the program while using the **Skip extraction** option.

These outputs are organized as follows:

```
Output directory                                 Required for running with
├── lipid_data.json                                  Skip extraction option
└── Module1_Outputs
         ├── CombinedAnalysis.csv
         ├── meanAllSigma.csv
         ├── Intersected_gauss.csv
         ├── extract_{isomer1}_MaxCov.csv
         ├── extract_{isomer2}_MaxCov.csv
         ├── norm_{isomer1}.csv
         ├── norm{isomer2}.csv
         ├── norm_{isomer1}_parameters.csv
         ├── norm{isomer2}_parameters.csv
         └── Plots
└── Module2_Outputs
         ├── iDMS_params.csv
         ├── iDMS_training_history.csv
         ├── TrainingSet.csv
         ├── PredictionResult.csv
         └── PredictedSetPlots
```

Note: Changing the names of the required files and sub-directories for the **Skip extraction** option will generate an error and exit the current run.

For more information on the iDMS analysis pipeline, please refer to the complete program documentation: https://github.com/Neurolipidomics/iDMS/tree/main
